# Rates of molecular and morphological evolution are uncoupled across the eukaryote Tree of Life

**DOI:** 10.64898/2026.11.04.764458

**Authors:** Yasmin Asar, E. Karen López-Estrada, Manuela Cascini, Hervé Sauquet, Simon Y. W. Ho

## Abstract

A vast degree of phenotypic variation can be seen across lineages of eukaryotes. Although at least part of this diversity has a genomic basis, it is unclear whether there are broad-scale links that can be detected between rates of genomic and phenotypic evolution. The presence of a correlation would imply the action of shared drivers, such as mutational or adaptive processes that occur on a genome- and organism-wide scale. We investigated this question by using a phylogenetic approach to test for correlations between molecular and morphological evolutionary rates. We analysed 60 data sets of combined nucleotide sequences and discrete morphological characters, representing a diverse range of eukaryotes. Using a maximum-likelihood phylogenetic approach based on sister-pair contrasts, we found very little evidence of any link between rates of molecular and morphological evolution, a striking result given the scale and taxonomic breadth of our analyses. Our results point to contrasting evolutionary dynamics underlying genomes and phenotypic traits; these have led to greater variation in morphological evolutionary rates across lineages, whereas molecular evolution is more clocklike. Our study reveals the discrepancies in evolutionary drivers that act at the genomic and phenotypic levels across the eukaryotic Tree of Life.

## Introduction

The evolutionary process has generated an immense amount of genomic and phenotypic diversity across the Tree of Life, with organisms displaying a vast range of genome sizes and structures, body plans, and life histories. These patterns naturally raise the question of whether there is a strong relationship between genomic and phenotypic change, and more specifically whether rates of molecular and morphological evolution are connected. Genomic changes have a wide distribution of fitness effects and many nucleotide substitutions are likely to be neutral or nearly neutral (Kimura 1968; Ohta 1973), whereas morphological traits and other aspects of phenotype are often under selection (Darwin 1859). Furthermore, the heritability of phenotype can be highly complex, with individual morphological characters and other organismal traits potentially depending on hundreds to thousands of loci with small effects (Boyle et al. 2017). In contrast, some traits are governed by just one or a few loci (e.g., Antonarakis and Beckmann 2006), or by changes to regulatory elements and homeotic genes, which can drastically alter anatomical structures (Carroll 1995; Wray 2007). As a consequence, many researchers have concluded that rates of molecular and morphological evolution are only weakly coupled, if at all (e.g., Simpson 1953; Kimura 1983; Ohta 1992; Gould 2002).

Even in the apparent absence of direct links, molecular and morphological evolutionary rates might nevertheless display similar patterns of variation if they are subject to a shared driver. Some life-history characteristics and abiotic factors, such as generation length, body size, population size, and environmental stress, can conceivably affect rates of both molecular and morphological evolution (Omland 1997; Barraclough and Savolainen 2001). In particular, population size governs the efficacy of selection, such that the stochastic loss and fixation of genomic and morphological variants by drift will be a dominant mode of evolution in small populations (Ohta 1973). Thus, founder events during speciation would potentially cause rapid molecular evolution as well as accelerated morphological change (Mayr 1954; Eldredge and Gould 1972). During adaptive radiations, ecological opportunities might promote evolutionary innovations, generating conditions conducive to the joint elevation of molecular and morphological rates (Lee et al. 2013). Some macroevolutionary models even posit the existence of mechanisms that act jointly on rates of lineage diversification, molecular evolution, and morphological change (Budd and Mann 2025).

There have been several attempts to test the link between molecular and morphological evolutionary rates, with previous studies yielding conflicting results. For example, positive associations were found in seven of eight data sets from various animals and plants (Omland 1997) and in angiosperms (Barraclough and Savolainen 2001; Davies and Savolainen 2006), while arthropod lineages experienced elevated rates of molecular and morphological evolution during their early diversification in the Cambrian (Lee et al. 2013). However, subsequent reanalyses yielded mixed results, with some suggesting that methodological artefacts have had a confounding effect (Bromham et al. 2002) while others have confirmed a positive correlation between molecular and morphological rates (Seligmann 2010). Very weak or no associations between molecular and morphological evolutionary rates were found in studies of echinoids (Smith et al. 1992), island endemic taxa (Bromham and Woolfit 2004), mammals (Janecka et al. 2012), hymenopteran and hemipteran insects and seed plants (Barba-Montoya et al. 2021), angiosperms (Asar et al. 2023), stomatopods (Van Der Wal et al. 2025), and various eukaryotic taxa regarded as ‘living fossils’ (Cascini et al. 2026). Simulation-based evaluations have demonstrated that some methods for testing for correlated rates either lack power or have high false-positive rates, with their performance being dependent on tree size and other properties of the data (Asar et al. 2023). However, these evaluations also showed that some methods performed well under a range of realistic conditions, thus highlighting their potential for producing insights into the dynamics of evolutionary rates at the genomic and phenotypic scales.

Here we use a statistical phylogenetic approach on a large and comprehensive collection of data sets to test for associations between rates of molecular and morphological evolution. We characterize and compare these evolutionary rates for various groups across the eukaryote Tree of Life, using data sets from lineages including protists, seedless and seeded plants, marine invertebrates, arachnids, insects, fishes, and tetrapods. By pairing molecular and morphological data for the same taxa, we are able to draw direct comparisons between their evolutionary dynamics. Our study identifies greater variation in rates of morphological evolution than in rates of molecular evolution across lineages, while also revealing that these rates are decoupled throughout major lineages of eukaryotes.

## Results and Discussion

### Molecular and morphological evolutionary rates

Our analyses of 60 data sets, sampled from a diverse array of eukaryotic taxa, found widespread uncoupling of molecular and morphological evolutionary rates (Figure 1). These data sets had an average size of 24,564 bp and 301 morphological characters, ranging up to 394,684 bp and 4541 morphological characters (Supplementary Table S1). The number of sampled taxa in each data set ranged from 12 to 412, with the average tree size being 71 taxa. The data spanned a variety of genomic marker types and phenotypic features. For instance, we included nucleotide sequences from ultraconserved elements, protein-coding genes, and ribosomal RNA genes. The inclusion of nuclear, mitochondrial, and chloroplast markers allowed us to examine all available genomic compartments. Morphological data included skeletal, soft-tissue, external morphological, vegetative, and reproductive characters. Moreover, we aimed to sample as widely as possible from the eukaryote Tree of Life, encompassing unicellular organisms, clades of vascular and non-vascular land plants, as well as marine and terrestrial invertebrates and vertebrates. Although tetrapods were overrepresented among the 60 data sets, our sample also included representatives of major plant lineages and several protostome phyla.

**Figure 1.**
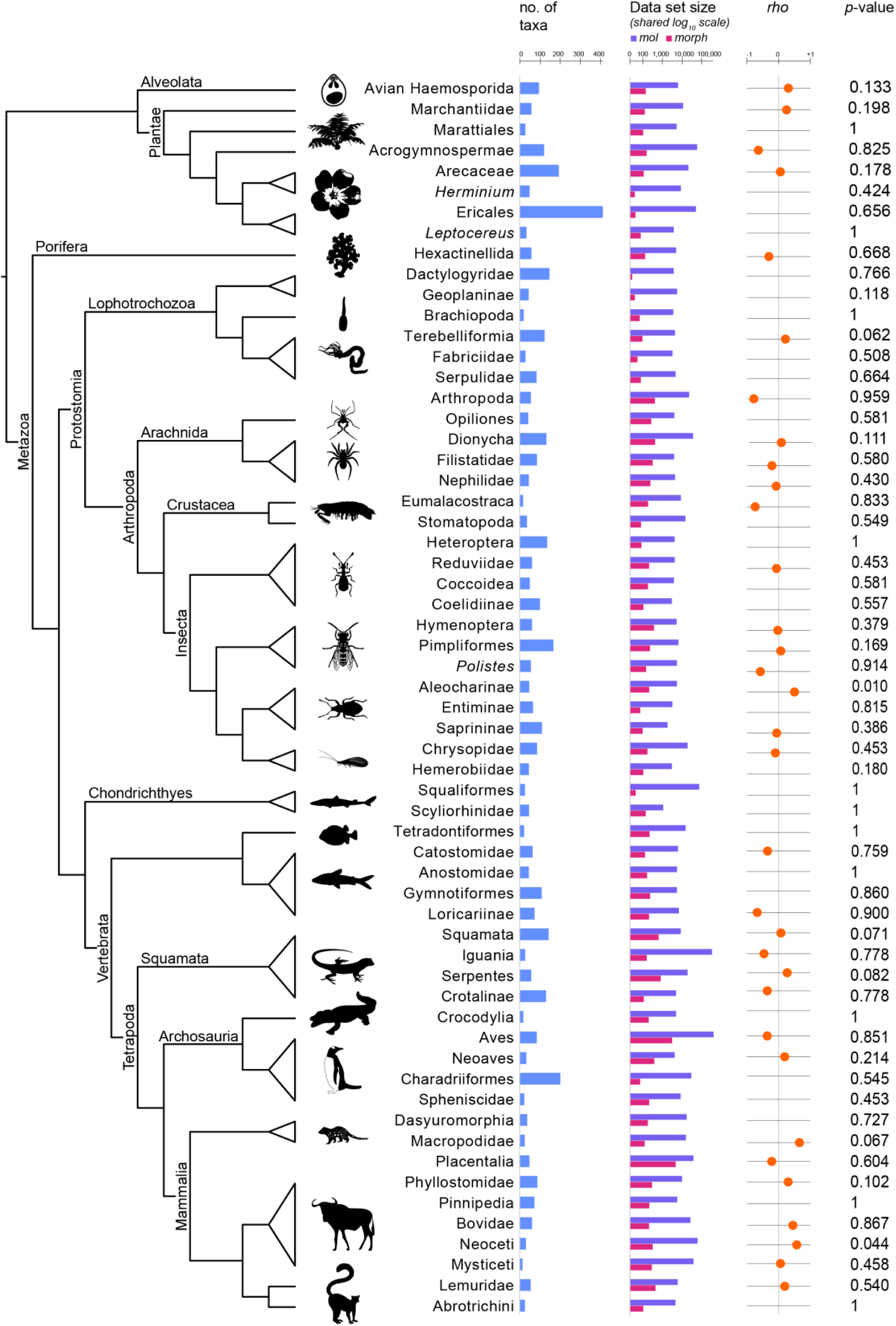
Tests of correlations between rates of molecular and morphological evolution in 60 data sets from eukaryotes. Tree shows evolutionary relationships among the organisms in the 60 data sets, with major clades labelled. Data columns show number of taxa (blue), sizes of molecular data sets (bp, purple) and morphological data sets (characters, dark pink), Spearman correlation coefficient (*rho*, orange) for data sets that passed the assumption tests, and *p*-values (for Spearman correlation coefficient where applicable, otherwise for sign test).

In most cases, we found no evidence of positive correlations between molecular and morphological evolutionary rates using independent sister-pair contrasts based on maximum-likelihood trees (Figure 1, Figure 2, Supplementary Figure S1). This method was previously validated in a comprehensive simulation study, which demonstrated its efficacy across a range of conditions and relevant data sizes (Asar et al. 2023). We found a significant positive correlation between molecular and morphological rates only in the data sets from the rove beetle subfamily Aleocharinae (*p* = 0.010) and from whales in the mammalian group Neoceti (*p* = 0.044). These also yielded high correlation coefficients (*ρ* = 0.64 and 0.71, respectively), but statistical support for these correlations disappears if any correction for multiple testing is applied. Thus, our analyses show that there is no broad association between rates of molecular and morphological evolution, even though our approach is expected to have sufficient statistical power to identify any such relationships should they be present (Asar et al. 2023).

**Figure 2.**
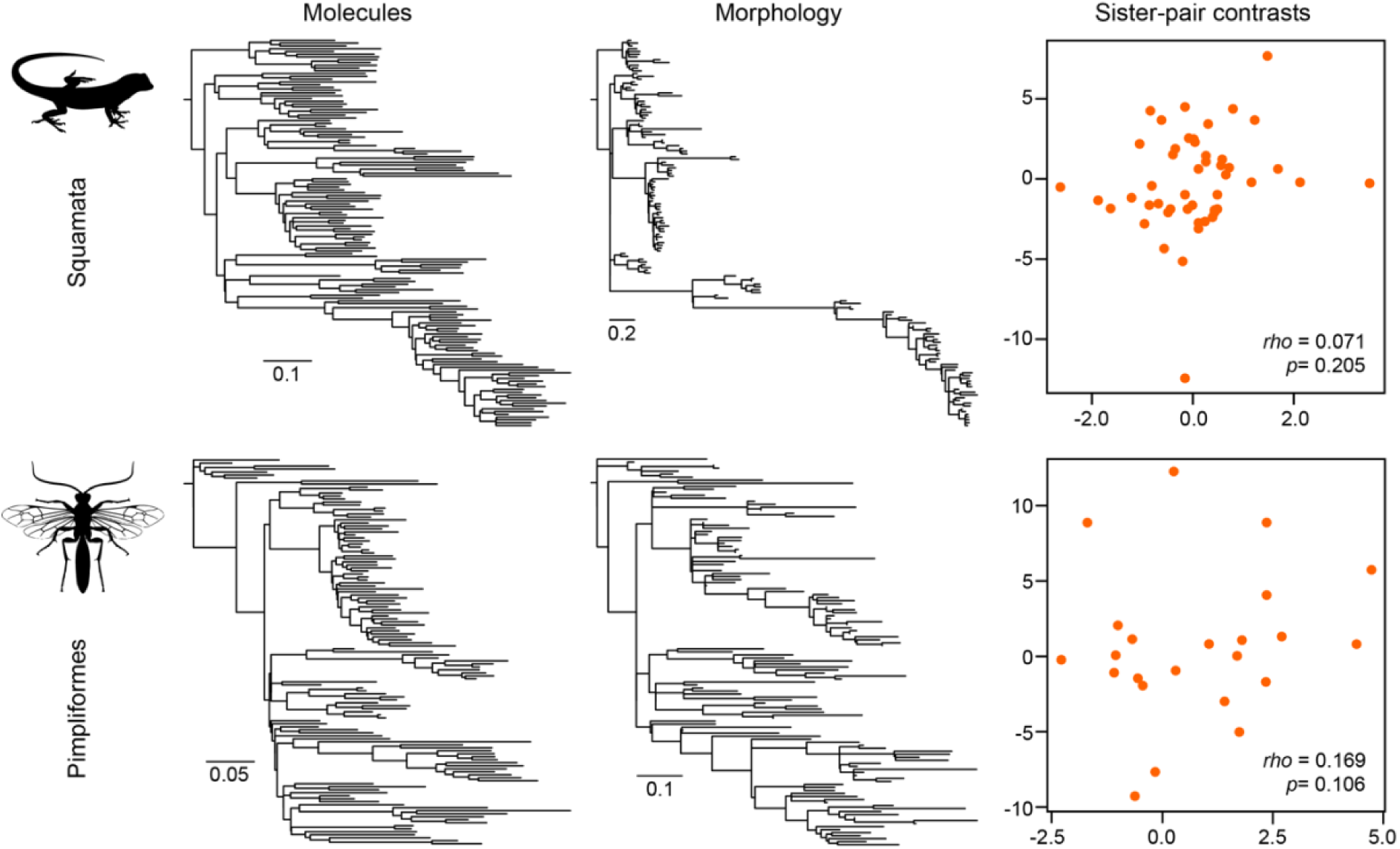
Tests of correlation between molecular and morphological evolutionary rates, shown for examples from lizards and snakes in the reptile order Squamata (top row) and parasitoid wasps in the group Pimpliformes (bottom row). These examples, which show a lack of correlation between molecular and morphological rates, are presented because they are based on large data sets: 8280 bp and 610 morphological characters from 141 taxa of Squamata (Gauthier et al. 2012; Pyron 2017), and 6213 bp and 222 morphological characters from 165 taxa of Pimpliformes (Spasojevic et al. 2021). Left column shows phylograms inferred from molecular sequence data, with scale bars denoting substitutions per site. Middle column shows phylograms inferred from morphological character data, with scale bars denoting changes per character. Right column shows plots of standardized, log-transformed sister-pair contrasts of molecular branch lengths and morphological branch lengths, with Spearman correlation coefficient (*rho*) and *p*-value calculated using one-sided Spearman’s rank correlation test. Plots for the 33 data sets that passed the assumption tests are presented in Supplementary Figure S1.

The results of our analyses stand in contrast with those from some of the earliest studies, which detected weak but significant positive associations between molecular and morphological rates in data sets from invertebrates, tetrapods, and flowering plants (Omland 1997; Barraclough and Savolainen 2001; Davies and Savolainen 2006). However, our finding of uncoupled molecular and morphological evolutionary rates is concordant with the results of several more recent studies (Janecka et al. 2012; Halliday et al. 2019; Barba-Montoya et al. 2021; Simões et al. 2020; Asar et al. 2023; Van Der Wal et al. 2025), while also confirming that this pattern occurs across a much broader set of taxa. One recent study of dozens of taxa regarded as ‘living fossils’ found a slowdown in morphological evolution in these lineages, but without being accompanied by substantial reductions in the rate of molecular evolution (Cascini et al. 2026); however, those taxa represent a biased sample of organisms that have apparently experienced some degree of morphological stasis. Thus, by analysing genomic sequences and morphological characters on a much larger scale than previous studies, we find no compelling evidence for a general link between rates of molecular and morphological evolution across the eukaryote tree.

Our failure to detect an association between molecular and morphological rates might have been partly due to shortcomings in the data and our methodological approach. Most of the data sets had at least 100 morphological characters, but these characters might have strong biases in sampling. For example, parsimony-informative characters are often sampled in preference to autapomorphies (Yeates 1992; Matzke and Irmis 2018), a collection bias that can lead to an apparent inflation of morphological evolutionary rates along internal branches compared with terminal branches (Seligmann 2010; Lee and Palci 2015). Moreover, the sister-pairs approach used in this study does not use information from all of the taxa in each data set. Greater statistical power can be achieved by testing for correlations using branch-rate estimates from a molecular dating analysis, as done in several previous studies (Asar et al. 2023; Asar et al. 2024; Van Der Wal et al. 2025). However, our preliminary analyses using such an approach revealed that many estimates of morphological branch rates had high uncertainty, with very wide 95% credible intervals, so we chose to take a conservative approach and limit our investigations to the sister-pairs contrasts. In a recent study, the sister-pairs approach was able to detect correlations in rates between nuclear, chloroplast, and mitochondrial genomes in multiple data sets from plants, producing results concordant with those from analyses of Bayesian branch rates (Asar et al. 2024). To achieve more powerful tests of correlations between molecular and morphological evolutionary rates, future studies would benefit from analysing using expanded morphological data sets, including more comprehensive sampling of morphological characters and taxa.

### Disparities in evolutionary rate variation across lineages

The uncoupling of molecular and morphological rates can be partly explained by the disparity in their levels of among-lineage variation, which we quantified by the coefficient of variation (CV) of root-to-tip distances in the phylogenetic trees (Figure 3, Supplementary Table S2). Values of CV close to 0 indicate strictly clocklike evolution, with higher values reflecting larger departures from rate constancy. We found significantly lower levels of among-lineage rate heterogeneity in the molecular data than in the morphological data, with respective mean CVs of 0.25 and 0.39 (Wilcoxon signed-rank test *p* = 5.02×10^-8^). These results were qualitatively consistent with estimates of CVs of branch rates that we obtained using a Bayesian phylogenetic approach, but which we do not report here because of the imprecision in branch-rate estimates obtained using that approach (as described in the preceding section).

**Figure 3.**
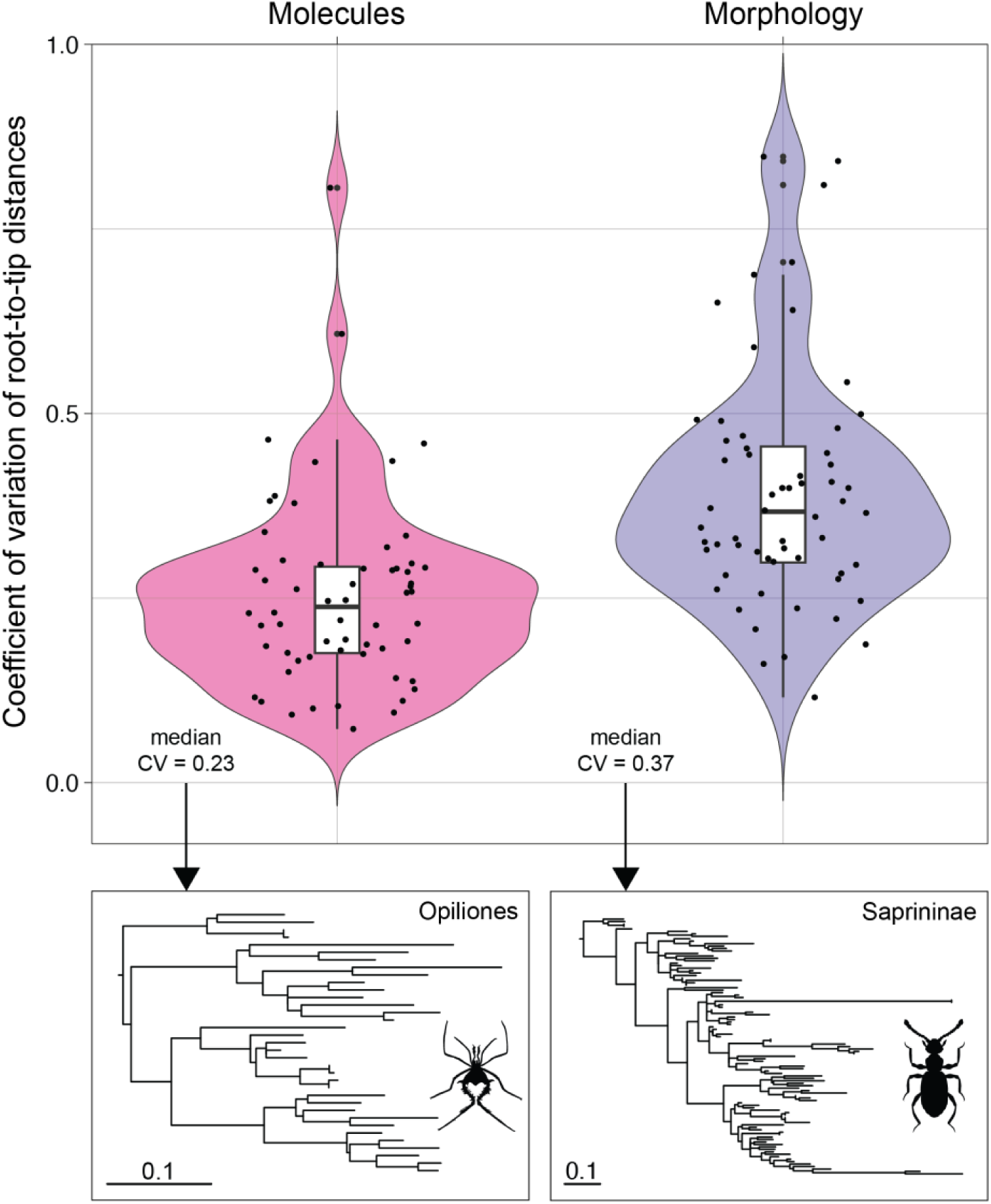
Evolutionary rate variation across lineages, shown as overlaid violin plots and boxplots of the coefficient of variation (CV) of root-to-tip distances for molecular sequence alignments (*n* = 60) and morphological character matrices (*n* = 60). For each data set, the CV value was calculated as the standard deviation in root-to-tip distances divided by the mean root-to-tip distance in the maximum-likelihood phylograms. Phylograms are shown for the molecular data set from harvestmen in the arachnid order Opiliones and the morphological data set from the clown beetle subfamily Saprininae, which yielded the median CVs shown in the main plot.

Many of the morphological data sets in our analyses exhibited large degrees of rate heterogeneity across lineages. Their CVs ranged from 0.12 (crevice weaver spiders in the arachnid family Filistatidae) to 0.85 (lizards and snakes in the reptile order Squamata), with CV > 0.30 for 44 of 60 data sets (Supplementary Table S2). Our results suggest that rate variation is a common feature of morphological evolution across eukaryotes, confirming previous findings of morphological rate variability based on different approaches (Van Valen 1974; Gingerich 1983; Hunt 2007; Rabosky et al. 2013; Gingerich 2019) and on phylogenetic studies of particular groups of organisms (e.g., Beck and Lee 2014; O’Reilly et al. 2015; Lee 2016; Parins-Fukuchi and Brown 2017; Klopfstein et al. 2019). Many morphological and other phenotypic traits are likely to be subject to selection pressures (Darwin 1859), with perhaps only a small proportion evolving neutrally (Ho et al. 2017; Zhang 2018). In general, we would expect the mode, direction, and strength of selection to be uneven across taxa, leading to variable rates of evolutionary change across lineages and across suites of characters (Lynch 1990; Hoekstra et al. 2001). Our results are consistent with previous claims that evolutionary processes generate much more heterogeneous patterns of morphological character variation than molecular sequence variation (Goloboff et al. 2019).

The low among-lineage rate heterogeneity that we found in most molecular data sets (CV < 0.30 for 47 of 60 data sets; Supplementary Table S2) indicates that molecular evolution is generally more clocklike than morphological evolution. Some data sets showed exceptionally low rate variation across lineages, with CVs below 0.1; these included the carnivorous marsupials in the order Dasyuromorphia, leafhoppers in the bug family

Coelidiinae, and kangaroos, wallabies, and relatives in the marsupial family Macropodidae. Although there are some instances of high molecular rate variation across lineages, as seen in our results from the crustacean subclass Eumalacostraca and fish order Tetraodontiformes, we also found instances of low among-lineage rate variation even in data sets with phylogenetically diverse sampling (Supplementary Table S2). These patterns broadly suggest a constant accumulation of genetic change over geological time, consistent with the general predictions of neutral and nearly neutral theories of molecular evolution (Kimura 1968; Ohta 1973). Although previous work has revealed that variation in molecular evolutionary rates is ubiquitous (Ho 2020), with links to generation length, diversification rates, and other factors (Bromham 2020), our results indicate that the magnitude of this variation is generally modest in most groups of eukaryotic organisms.

We detected no association between the CVs for molecular and morphological rates (*p* = 0.084, Spearman’s rank correlation test). For 48 of 60 data sets, CVs were smaller for molecular data than for morphological data. In many groups of taxa, we found rate constancy in molecular evolution contrasting with substantial levels of rate heterogeneity in morphological evolution (Supplementary Table S2). For example, armoured catfishes in the subfamily Loricariinae had molecular rates with CV = 0.17 and morphological rates with CV = 0.84, whereas monogenean flatworms in the family Dactylogyridae had molecular rates with CV = 0.21 and morphological rates with CV = 0.81. Our results thus provide compelling evidence that patterns of among-lineage rate variation differ between molecular and morphological evolution, and that there is greater heterogeneity in morphological rates of change.

### Separate drivers of molecular and morphological evolution

Our analyses have shown that molecular and morphological rates do not vary in concert across lineages, suggesting that distinct evolutionary dynamics predominate at the genomic and phenotypic levels. However, we note that our study is based on data at the lowest genomic level (primary nucleotide sequence) and at one of the highest phenotypic levels (morphological characters). Selection is likely to be important across multiple phenotypic levels, but any changes in morphological traits might only have a very weak association with nucleotide substitutions in the genome. Correspondences between coding sequences and phenotypic traits can be masked by epistasis, pleiotropy, genetic redundancy, and gene-by-environment interactions (de Visser et al. 2003; Hansen 2006; Dowell et al. 2010).

Furthermore, the evolution of morphological traits can be driven by other forms of genomic change or alterations in regulatory potential (Irimia and Maeso 2019; Recuerda and Campagna 2024). For example, macroevolutionary change can be driven by changes in gene content, perhaps through changes in repeats, whole-genome duplications, or gene losses and gains (e.g., Fondon and Garner 2004; Clark and Donoghue 2018; Sharma et al. 2018), or by expansion of homeotic gene clusters (Carroll 1995; Lemons and McGinnis 2006).

Our study predominantly deals with nucleotide sequences from coding regions, because these data are the most widely accessible and available. Many of the data sets analysed in this study comprise commonly used genetic markers from the mitochondrial and nuclear genomes, including genes without known associations with diagnosable morphological variation. Across the genome, only a small proportion of nonsynonymous nucleotide substitutions and even fewer synonymous substitutions are likely be adaptive (Eyre-Walker and Keightley 2007). However, substitution rates in a sufficiently representative set of nucleotide sequences can still provide a useful proxy for the genome-wide rate of molecular evolution in the taxa being analysed. Our results thus show that higher rates of molecular evolution do not necessarily translate into higher rates of morphological change, although this general pattern does not exclude the possibility of correlations in rates between specific genes and subsets of phenotypic traits (e.g., Barrier et al. 2001). Further investigation of genomic structure, regulatory regions, and conserved non-coding elements will provide a deeper understanding of rates of genomic evolution and their relationship to phenotypic change. Such studies will become increasingly feasible with the rapid growth of high-quality reference genomes (e.g., Formenti et al. 2026).

Some models of macroevolution propose a punctuated mode of morphological change, with varying statements about the association with molecular change. For instance, morphological evolution might be propelled by novel ecological opportunities during adaptive radiations (Simpson 1953), occur in discrete pulses over time (Uyeda et al. 2011; Landis and Schraiber 2017), or experience shifts in directionality (Pagel et al. 2022). Under the punctuated equilibrium model, bursts of morphological evolution take place at speciation events, but phenotypic change is otherwise characterised by stasis (Eldredge and Gould 1972). The punctuated equilibrium model has been extended to the genome level, where some have argued that bursts of genetic change coincide with speciation (Nichol et al. 1993; Venditti and Pagel 2010). This concept is prominently expressed in the proposal by Mayr (1954) that speciation is tied to genetic revolutions that occur during founder events. Bursts of genetic change have been detected at divergence events in specific groups of organisms, such as snakes (Manceau et al. 2020) and Australian monitor lizards in the family Varanidae (Janzen et al. 2022), and in broader studies of animals (Yuan et al. 2026) and eukaryotes (Pagel et al. 2006). Nevertheless, there has been little evidence of punctuations of genomic and phenotypic change occurring in synchrony (López-Estrada et al. 2025). Further insight into these patterns of evolutionary change can potentially be gained by using recently developed models for detecting and quantifying pulses of evolution and shifts in macroevolutonary mode (Manceau et al. 2020; Janzen et al. 2022; Pagel et al. 2022; Douglas et al. 2025; Wright and Wagner 2025).

### Implications for evolutionary analyses

The disjunction between molecular and morphological rates has important implications for evolutionary analyses of combined data sets, including phylogenetic inference and molecular dating. When the data comprise combinations of heterogeneous data types or different suites of phenotypic characters, measures must be taken to account for their disparate patterns of among-lineage rate heterogeneity (Clarke and Middleton 2008; Lee 2016; Goloboff et al. 2019). Our analyses revealed that large disparities in the pattern and degree of among-lineage rate variation between molecular and morphological data are commonplace. In these cases, the data set needs to be partitioned for phylogenetic analyses, with independent sets of branch lengths enabled for each subset of the data. In the context of molecular dating, including total-evidence dating analyses that rely on morphological data from extant and fossil taxa (Ronquist et al. 2012; Álvarez-Carretero et al. 2019), a separate relaxed clock might be needed for each data subset in order to accommodate the complex patterns of rate variation (Duchêne et al. 2016). These forms of partitioning would potentially lead to a substantial increase in model complexity, especially when compared with simpler models involving proportionate branch lengths or linked clock models. Accordingly, the optimal scheme should be identified using a model-selection approach (e.g., Duchêne et al. 2016; Duchêne et al. 2020).

The among-lineage variation that we identified in morphological evolutionary rates underscores the difficulty in phylogenetic dating using morphological data. Although there are growing numbers of studies that have inferred evolutionary timescales using morphological data alone (e.g., Lee et al. 2014) or in combination with molecular data (e.g., Gavryushkina et al. 2017; Cascini et al. 2019), high levels of among-lineage rate variation present considerable challenges to the performance of phylogenetic methods and clock models (Lyons-Weiler and Takahashi 1999; Lee and Palci 2015; Vankan et al. 2022). In cases of extreme rate variation, phylogenetic analyses face the risk of clock-model misspecification or even model inadequacy (Duchêne et al. 2015). Models of macroevolution, such as those that account for punctuated equilibrium or other bursts of evolutionary change (e.g., Manceau et al. 2020; Douglas et al. 2025), might encounter difficulty in resolving the contributions of these spikes in the presence of pronounced among-lineage rate heterogeneity. Our results call for careful evaluation of rate variation during the course of phylogenetic inference, along with rigorous selection of models in evolutionary analyses.

## Conclusions

Our study has revealed that rates of molecular and morphological evolution are uncoupled across a diverse range of eukaryotic groups. In general, molecular evolution appears to be more clocklike than morphological evolution, although examples of extreme rate variation can be found in both data types. Our results are consistent with disparate mechanisms of evolution operating at the genomic and phenotypic scales, lending some support to the neutral theory of molecular evolution and consistent with a dominant role for adaptive evolution in morphological traits. Further insights into the underlying mechanisms of genomic and phenotypic change can be gained through focused studies of genomic components such as homeotic genes and regulatory regions, comparisons of genomic markers, and analyses of comprehensive morphological and phenotypic data sets.

## Materials and methods

### Data sets

To allow comparison of the evolutionary rates between molecular sequences and morphological characters, we assembled a collection of 60 combined data sets from a diversity of eukaryotic groups (Figure 1, Supplementary Table S1). We prioritized data from total-evidence dating studies, to maximize the amount of molecular and morphological data from matching taxa. When we obtained data sets that were from different source publications, we pruned the data sets to leave only the intersecting taxa. Thus, fossil taxa were automatically excluded from our analyses owing to the lack of matching molecular data. We aimed to include molecular data sets that sampled as many nuclear, mitochondrial, and/or chloroplast markers as possible. Similarly, we aimed to include morphological data sets that sampled at least 100 discrete morphological characters. Our resulting data sets generally had a minimum of 50 matching taxa, to ensure sufficient statistical power for our analyses (Asar et al. 2023). However, we waived this condition for taxon-poor groups such as brachiopods, pinnipeds, and crocodilians, as long as there were at least 10 ingroup taxa. For our analyses of morphological data, we removed constant characters and treated polymorphic states (e.g., coded as “0/1/2”) as missing data, so they were not used in our estimation of morphological rates. Overall, the mean sequence alignment length used in analyses was 24,564 bp, the mean number of discrete morphological characters in the matrices was 301, and the mean number of intersecting taxa was 71 (Figure 1, Supplementary Table S1).

To enable comparison of rates of molecular and morphological evolution in a phylogenetic framework, we required a constraint topology for analyses. We inferred the constraint topology by analysing combined molecular and morphological data in IQ-TREE2 (version 2.2.2.6; Bui et al. 2020). The molecular data and morphological data (each in their own subset with proportionate branch lengths) were analysed using IQ-TREE2 with 1000 bootstrap replicates. The GTR+Γ model was used for the molecular data and the MK+ASC+Γ model was used for the morphological data (Lewis 2001). For the sake of consistency, we used the combined data to infer the constraint tree for each group of taxa in our study. These trees were generally consistent with those reported in the original studies describing these data sets (Supplementary Table S1).

### Phylogenetic analyses

We first describe the partitioning scheme and models of evolution used for our maximum-likelihood analyses of molecular and morphological data sets. For molecular data, individual loci were loaded as separate subsets, with selection of the partitioning scheme and substitution models conducted using IQ-TREE2. For the morphological data, the discrete characters were simply partitioned according to their number of possible states. This partitioning scheme was chosen to follow standard treatments in software used to analyse morphological characters. For each subset of the morphological data, we used the Mk model with ascertainment-bias correction and a discrete gamma distribution to model rate heterogeneity across sites (MK+ASC+Γ). We then estimated branch lengths for the molecular and morphological data sets using maximum likelihood in IQ-TREE2 (all phylograms are presented in Supplementary Figure S1). This was performed in separate analyses for molecular and morphological data sets, with a constrained topology, models, and data partitioning as described above.

For each data set, we estimated branch rates using Bayesian inference in BEAST2 (version 2.7.7; Bouckaert et al. 2019). These estimates can enable powerful tests of correlations between molecular and morphological rates, even for relatively small data sets (Asar et al. 2023). We followed the same partitioning schemes used for the maximum-likelihood analyses described above, while using bModelTest (Bouckaert and Drummond 2017) to average the substitution model for each molecular subset and using the MK+ASC+Γ model for each morphological subset. Separate uncorrelated lognormal relaxed clocks for specified for the molecular and morphological data (Drummond et al. 2006), with the mean rate of the morphological clock fixed to 1.0. For a large proportion of data sets, the estimates of morphological branch rates were highly uncertain, with wide 95% credible intervals. Consequently, we chose not to proceed with further analyses of these branch-rate estimates.

### Independent sister-pair contrasts

We tested for correlations between phylogenetically independent sister-pair contrasts using the molecular and morphological phylograms obtained by maximum-likelihood analysis. For each phylogram, independent sister-pairs were obtained using the R package *diverge* (version 2.0.6; Anderson and Weir 2020). Each sister-pair shares a most recent common ancestor to the exclusion of all other taxa in the tree, so that each member of the pair has had the same amount of time for evolutionary change to accumulate. Thus, any difference in the branch lengths leading to the two members of each sister-pair can be considered as a difference in the rate of evolution (Lanfear et al. 2010), and these contrasts can be compared between molecular and morphological phylograms to determine whether there is any correlation.

We tested the assumption of homogeneity of variances for the independent contrasts using the tests recommended previously (Garland et al. 1992; Freckleton 2000; Welch and Waxman 2008). We started by checking for an association between the standardized contrasts (absolute values) and their standardization factors (Assumption Test 1). The standardization factors were calculated by taking the square root of the sum of the branch lengths for each contrast. This test also allowed us to visualize and remove any shallow contrasts that were unusually influential (Garland et al. 1992, Welch and Waxman 2008). If there is an association between the standardized contrasts and their standardization factors, even after removing unsuitable contrasts, then only a two-sample sign test should be used to evaluate correlations between independent contrasts (Garland et al. 1992, Welch and Waxman 2008). A two-sample sign test was used for 27 of 60 data sets (Figure 1), due to violations of Assumption Test 1.

The second assumption test was to check for an association between the mean sum of the branch lengths and the contrasts (absolute values) (Assumption Test 2). Any association between the mean sum and contrasts indicates that a non-parametric test should be used to assess correlations between molecular and morphological independent sister-pair contrasts (Freckleton 2000, Welch and Waxman 2008). There were a few instances in which Assumption Test 2 was violated, so we implemented a non-parametric Spearman’s rank correlation test for all analyses of independent sister-pair contrasts.

All contrasts were log-transformed for assumption tests. The results of assumption tests and the number of contrasts used in analyses are provided in Supplementary Table S3. The log-transformed contrasts were standardized using the square root of the sum of the branch lengths for each contrast, following the recommendation of Welch and Waxman (2008). We used Spearman’s rank correlation test to check for positive associations between molecular and morphological contrasts.

## Acknowledgements

The authors acknowledge the technical assistance provided by the Sydney Informatics Hub, a Core Research Facility of the University of Sydney.

