## Supplementary Material for "Rates of molecular and morphological evolution are uncoupled across the eukaryote Tree of Life"


| | Page |
|---|---|
| **Supplementary Table S1.** Details of 60 molecular and morphological data sets analysed in the study. | 2 |
| **Supplementary Table S2.** Coefficient of variation (CoV) of root-to-tip distances estimated for molecular and morphological data sets. | 13 |
| **Supplementary Table S3.** Details of assumption tests and number of contrasts used for independent sister-pair contrast analyses. | 15 |
| **Supplementary Figure S1.** Plots of standardized, log-transformed sister-pair contrasts of molecular branch lengths and morphological branch lengths. | 17 |
| **References** | 18 |

**Supplementary Table S1.** Details of 60 molecular and morphological data sets analysed in the study. Many of the listed source publications presented modified versions of morphological data sets from earlier studies, but for the sake of brevity we have only cited the publications that presented the version of the data set that we used in our analyses.

| **Clade** | **Taxon** | **Molecular data**[b] | **Morphological data**[c] | **Inter-secting taxa**[d] | **No. of sister pairs** | **Source** |
|---|---|---|---|---|---|---|
| Alveolata | Avian Haemosporida | 5991 bp from 93 taxa; 3 mt genes *COX1*, *COX3*, *CYTB* | 133 chars from 93 taxa; blood stages, parasite development | 93 | 26 | Hernández-Lara et al. (2018) |
| Plantae: Marchantiophyta | Marchantiidae | 10,977 bp from 62 taxa; 1 nuc, 7 cp, 3 mt genes | 120 chars from 62 taxa; gametophytic and sporophytic phase, cytological, developmental, sexual, and macroscopic characters | 56 | 13 | Flores et al. (2020) |
| Plantae: Monilophyta | Marattiales | 5104 bp from 33 taxa; 4 cp markers (*atpB*, *rbcL*, *rps4* and *rps4*-*trnS* spacer, *trnS*-*trnG* spacer and *trnG* intron) | 98 chars from 89 taxa; anatomy and morphology | 26 | 5 | May et al. (2021) |

| | | | | | | |
|---|---|---|---|---|---|---|
| Plantae: Spermatophyta | Acrogymnospermae | 57,015 bp from 1055 taxa; 78 cp genes | 148 chars from 137 taxa; life-history, vegetative and reproductive characters | 119 | 39 | Stull et al. (2021) |
| Plantae: Monocotyledoneae | Arecaceae | 20,355 bp from 205 taxa; cp genes, low-copy nuc genes and ribosomal nuc DNA | 105 chars from 192 taxa; vegetative and reproductive characters | 192 | 59 | Baker et al. (2009) |
| Plantae: Monocotyledoneae | *Herminium* | 8303 bp from 47 taxa; 2 nuc (*ITS*, *Xdh*) and 5 cp markers (*matK*, *psaB*, *psbA-trnH*, *rbcL*, *trnL-F*) | 37 chars from 56 taxa; vegetative and reproductive characters | 47 | 14 | Raskoti et al. (2015) |
| Plantae: Eudicotyledoneae | Ericales | 49,435 bp from 4943 taxa; 25 cp, mt, and ribosomal markers | 40 chars from 415 taxa; floral characters | 412 | 126 | Rose et al. (2018); Sauquet (2019) and Herting et al. (2023) |
| Plantae: Eudicotyledoneae | *Leptocereus* | 3628 bp from 46 taxa; 5 cp markers, 4 intergenic spacers, and 1 intron | 74 chars from 46 taxa; vegetative and reproductive characters | 32 | 14 | Barrios et al. (2020) |
| Porifera | Hexactinellida[a] | 4806 bp from 73 taxa; nuc markers (*18S* and *28S*) and mt (*16S* and *COX1*) | 124 chars from 124 taxa; external morphology | 56 | 18 | Dohrmann et al. (2017) |
| Platyhelminthes: Monogenea | Dactylogyridae | 3627 bp from 154 taxa; 4 markers (*18S*, *28S*, *COX1*, *ITS1*) | 22 chars from 148 taxa; external morphology | 145 | 45 | Kmentová et al. (2022) |

| | | | | | | |
|---|---|---|---|---|---|---|
| Platyhelminthes: Tricladida | Geoplaninae | 5414 bp from 74 taxa; 6 markers (*18S*, *28S*, *COX1*, *EF*, *DOM5*, *Tnuc813*) | 37 chars from 47 taxa; body, sensory organs, gland cells, muscles, external morphology, reproductive parts | 42 | 15 | Lago-Barcia et al. (2023) |
| Brachiopoda | Brachiopoda | 3435 bp from 18 taxa; ribosomal DNA | 66 chars from 26 taxa; interior and exterior shell characters | 18 | 3 | Cohen & Bitner (2013); Bapst et al. (2018) |
| Annelida: Polychaeta | Terebelliformia | 4159 bp from 133 taxa; 5 markers (*16S*, *18S*, *28S*, *COX1*, *H3*) | 90 chars from 133 taxa; external morphology | 121 | 44 | Stiller et al. (2020) |
| Annelida: Polychaeta | Fabriciidae[a] | 3107 bp from 26 taxa; 3 nuc markers (*18S*, *28S*, *H3*) | 50 chars from 78 taxa; adult morphology; reproductive system | 26 | 9 | Huang et al. (2011) |
| Annelida: Polychaeta | Serpulidae | 4483 bp from 93 taxa; 4 markers (*18S*, *28S*, *CYTB*, *H3*) | 75 chars from 93 taxa; external morphology | 81 | 21 | Kupriyanova et al. (2023) |
| Arthropoda | Arthropoda | 22,131 bp from 53 taxa; 62 protein-coding genes | 395 chars from 53 taxa; diverse internal and external morphological traits | 53 | 16 | Lee et al. (2013) |
| Arthropoda: Arachnida | Opiliones[a] | 3885 bp from 63 taxa; *18S* and *28S* | 253 chars from 44 taxa; external and internal | 40 | 13 | Giribet et al. (2002); Garwood et al. (2011) |

| | | | | | | |
|---|---|---|---|---|---|---|
| | | | anatomy; reproductive system | | | |
| Arthropoda: Arachnida: Araneae | Dionycha | 35,106 bp from 156 taxa; 197 UCEs and 6 markers (*12S*, *16S*, *18S*, *28S*, *COX1*, *H3*) | 400 chars from 130 taxa; morphological and behavioural characters | 130 | 40 | Azevedo et al. (2022) |
| Arthropoda: Arachnida: Araneae | Filistatidae | 3787 bp from 133 taxa; 4 markers (*COX1*, *16S*, *H3*, *28S*) | 302 chars from 103 taxa; external morphology, genitalia, respiratory system, organs, internal features, and behavioural traits | 83 | 18 | Magalhaes & Ramírez (2022) |
| Arthropoda: Arachnida: Araneae | Nephilidae[a] | 4197 bp from 65 taxa; 8 mt and nuc markers | 231 chars from 61 taxa; morphological and behavioural characters | 43 | 11 | Kuntner et al. (2013) |
| Arthropoda: Crustacea | Eumalacostraca[a] | 8316 bp from 24 taxa; 4 mt and nuc markers (*16S*, *COX1*, *18S*, *28S*) | 177 chars from 24 taxa; external morphology; internal organs; reproductive and developmental features | 15 | 4 | Jenner et al. (2009) |
| Arthropoda: Crustacea | Stomatopoda | 14,432 bp from 34 taxa; whole mt genomes and 3 nuc markers | 77 chars from 34 taxa; external morphology | 34 | 11 | Van Der Wal et al. (2025) |

| | | | | | | |
|---|---|---|---|---|---|---|
| Arthropoda: Insecta: Hemiptera | Heteroptera | 4018 bp from 135 taxa; 3 genes *16S*, *18S*, *28S* | 81 chars from 135 taxa; external morphology | 134 | 43 | Weirauch et al. (2019) |
| Arthropoda: Insecta: Hemiptera | Reduviidae | 4055 bp from 59 taxa; 6 mt and nuc genes | 199 chars from 156 taxa; external morphology, genitalia, glands, sexual dimorphism | 59 | 19 | Forthman & Weirauch (2017) |
| Arthropoda: Insecta: Hemiptera | Coccoidea | 3731 bp from 51 taxa; partial nuc markers (*18S*, *28S*, *EF1-α* ) | 174 chars from 112 taxa; external morphology, genitalia | 48 | 13 | Vea & Grimaldi (2016); Vea & Grimaldi (2015) |
| Arthropoda: Insecta: Hemiptera | Coelidiinae | 2903 bp from 98 taxa; 2 mt genes (*COX1*, *16S*) and 4 nuc genes (*28S*, *H3*, *H2A*, *Wg*) | 102 chars from 98 taxa; external morphology, colouration, proportions, shape, texture, wing venation, genitalia | 98 | 28 | Wang et al. (2024) |
| Arthropoda: Insecta: Hymenoptera | Hymenoptera | 5096 bp from 66 taxa; 7 partial markers (*12S*, *16S*, *18S*, *28S*, *COX1*, *EF1-α*, *F2*) | 353 chars from 113 taxa; external morphology, genitalia | 59 | 17 | Ronquist et al. (2012) |
| Arthropoda: Insecta: Hymenoptera | Pimpliformes | 6213 bp from 207 taxa; 9 nuc protein-coding genes, partial *28S*, and mt *COX1* | 222 chars from 249 taxa | 165 | 41 | Spasojevic et al. (2021) |

| | | | | | | |
|---|---|---|---|---|---|---|
| Arthropoda: Insecta: Hymenoptera | *Polistes* | 5288 bp from 53 taxa; mt (*COX1*, *12S*, *16S*) and nuc genes (*18S*, *H3*, *EF1-* α) | 140 chars from 104 taxa; external morphology, male genitalia, larval traits, nest architecture | 53 | 11 | Santos et al. (2015); Somavilla et al. (2021) |
| Arthropoda: Insecta: Coleoptera | Aleocharinae | 5273 bp from 45 taxa; 9 markers (*18S*, *28S*, topoimerase I, *16S*, *ND1*, *Leu1*, *Leu2*, *COX1*, *COX2*) | 202 chars from 69 taxa; diverse characters sampled from across the external morphology | 45 | 14 | Orlov et al. (2021) |
| Arthropoda: Insecta: Coleoptera | Entiminae | 3084 bp from 67 taxa; mt genes (*16S*, *COX1*) and nuc genes (*18S*, *28S*) | 70 chars from 67 taxa; external morphology, genitalia | 64 | 18 | Marvaldi et al. (2018) |
| Arthropoda: Insecta: Coleoptera | Saprininae | 1752 bp from 112 taxa; mt *COX1* and nuc *18S* | 93 chars from 108 taxa; external morphology, genitalia | 108 | 35 | Lackner et al. (2024) |
| Arthropoda: Insecta: Neuroptera | Chrysopidae | 18,452 bp from 87 taxa; 21 mt and nuc genes | 165 chars from 177 taxa | 84 | 27 | Breitkreuz et al. (2022) |
| Arthropoda: Insecta: Neuroptera | Hemerobiidae | 2897 bp from 43 taxa; mt genes (*16S*, *COX1*) and nuc genes (*CAD*) | 101 chars from 43 taxa; external morphology, genitalia | 43 | 14 | Garzón-Orduña et al. (2016) |
| Fishes: Chondrichthyes | Squaliformes | 72,296 bp from 24 taxa; 172 exon sequences | 41 chars from 85 taxa; dental characters | 24 | 9 | Straube et al. (2015); Flammensbeck et al. (2018) |

| | | | | | | |
|---|---|---|---|---|---|---|
| Fishes: Chondrichthyes | Scyliorhinidae | 1044 bp from 44 taxa; mt *ND2* | 134 chars from 78 taxa; external morphology, musculature, denticles and osteology | 44 | 10 | Soares & Mathubara (2022) |
| Fishes: Osteichthyes | Tetraodontiformes[a] | 14,648 bp from 95 taxa; 16 mt and nuc genes | 210 chars from 56 taxa; internal and external osteological characters | 20 | 8 | Arcila et al. (2015) |
| Fishes: Osteichthyes | Catostomidae | 5925 bp from 85 taxa; mt genes (*CYTB*, *ND2*, *COX1*) and nuc genes (*IRBP*, *RPS7*, *GHI*) | 123 chars from 64 taxa; morphological and biochemical characters | 62 | 20 | Bagley et al. (2018) |
| Fishes: Osteichthyes | Anostomidae | 5321 bp from 55 taxa; mt genes (*COX1*, *CYTB*, *16S*) and nuc genes (*Myh6*, *RAG1*, *RAG2*) | 158 chars from 58 taxa; osteology | 43 | 11 | Sidlauskas et al. (2022) |
| Fishes: Osteichthyes | Gymnotiformes | 5277 bp from 149 taxa; mt genes (*16S*, *COX1*, *CYTB*) and nuc genes (*ZIC*, *RAG1*, *RAG2*) | 223 chars from 166 taxa; osteology, musculature, colouration, transformed quantitative traits, neurology | 107 | 32 | Tagliacollo et al. (2016) |
| Fishes: Osteichthyes | Loricariinae | 6623 bp from 86 taxa; mt genes (*CYTB*, *ND2*, *12S*, | 196 chars from 90 taxa; osteology and external morphology | 71 | 22 | Londoño-Burbano & Reis (2021) |

| | | | | | | |
|---|---|---|---|---|---|---|
| | | *16S*) and nuc genes (*Myh6*, *RAG1*, *RAG2*) | | | | |
| Reptilia: Squamata | Squamata | 8280 bp from 141 taxa; 3 mt genes (*12S*, *16S*, *CYTB*) and 3 nuc genes (*BDNF*, *CMOS*, *RAG1*) | 610 chars from 192 taxa; diverse morphological characters | 141 | 53 | Pyron (2017); Gauthier et al. (2012) |
| Reptilia: Squamata | Iguania | 326,304 bp from 44 taxa; 839 UCEs | 152 chars from 31 taxa | 25 | 7 | Streicher et al. (2016); Smith (2009) |
| Reptilia: Squamata | Serpentes | 18,320 bp from 56 taxa; 1 mt and 21 nuc genes | 777 chars from 73 taxa; diverse morphological characters | 55 | 13 | Hsiang et al. (2015) |
| Reptilia: Squamata | Crotalinae | 4724 bp from 128 taxa; 5 mt genes (*CYTB*, *ND4*, *COX1*, *12S*, *16S*) and 4 nuc genes (*bdnf*, *c-mos*, *nt-3*, *rag-1*) | 107 chars from 129 taxa; external morphology | 128 | 33 | Carrasco et al. (2023) |
| Reptilia: Archosauria: Crocodylia | Crocodylia | 9284 bp from 16 taxa; mt and nuc markers | 189 chars from 16 taxa; osteology | 16 | 6 | Darlim et al. (2022) |
| Reptilia: Archosauria: Aves | Aves | 394,684 bp from 200 taxa; 259 nuc loci | 2954 chars from 188 taxa; largely osteology and myology | 82 | 24 | Prum et al. (2015); Livezey & Zusi (2006) |

| | | | | | | |
|---|---|---|---|---|---|---|
| Reptilia: Archosauria: Aves | Neoaves | 4038 bp from 32 taxa; nuc loci (*RAG1*, *RAG2*) | 368 chars from 40 taxa; osteology | 31 | 9 | Musser & Cracraft (2019) |
| Reptilia: Archosauria: Aves | Charadriiformes | 28,531 bp from 335 taxa; 15 mt and 12 nuc genes | 69 chars from 218 taxa; osteology | 199 | 68 | Černý & Natale (2022) |
| Reptilia: Archosauria: Aves | Spheniscidae | 8145 bp from 19 taxa; mt genes (*12S*, *16S*, *COX1*, *CYTB*) and nuc gene (*RAG1*) | 202 chars from 25 taxa; osteology, myology, integument, and reproductive biology | 21 | 7 | Gavryushkina et al. (2017) |
| Mammalia: Metatheria | Dasyuromorphia | 16,441 bp from 78 taxa; 7 mt genes and 8 nuc protein-coding genes or gene fragments | 173 chars from 54 taxa; postcranial and craniodental characters | 35 | 8 | Kealy & Beck (2017) |
| Mammalia: Metatheria | Macropodidae | 15,253 bp from 28 taxa; mt genomes | 119 chars from 28 taxa; craniodental and postcranial characters | 23 | 6 | Cascini et al. (2018) |
| Mammalia: Eutheria | Placentalia | 36,860 bp from 46 taxa; 27 nuc loci | 4541 chars from 86 taxa; cranial, dental, postcranial, soft-tissue characters | 46 | 14 | O'Leary et al. (2013) |
| Mammalia: Eutheria: Laurasiatheria | Phyllostomidae[a] | 9584 bp from 85 taxa; 10 mt and nuc genes | 278 chars from 113 taxa; dental characters | 85 | 24 | Dávalos et al. (2014) |

| | | | | | | |
|---|---|---|---|---|---|---|
| Mammalia: Eutheria: Laurasiatheria | Pinnipedia | 5490 bp from 22 taxa; 5 nuc protein-coding genes (*APOB*, *BRCA-1*, *RAG-1*, *IRBP*, *VWF*) | 204 chars from 66 taxa; craniomandibular, dental, postcranial characters | 22 | 4 | Paterson et al. (2020) |
| Mammalia: Eutheria: Laurasiatheria | Bovidae | 25,441 bp from 156 taxa; 9 nuc loci and complete mt genomes | 196 chars from 88 taxa; craniodental and postcranial characters | 61 | 19 | Calamari (2021) |
| Mammalia: Eutheria: Laurasiatheria | Neoceti[a] | 59,811 bp from 29 taxa; mt and nuc loci | 304 chars from 74 taxa; skeletal and dental morphology; digestive and sexual characters | 29 | 7 | Geisler et al. (2011) |
| Mammalia: Eutheria: Laurasiatheria | Mysticeti[a] | 37,013 bp from 90 taxa; mt and nuc loci | 272 chars from 90 taxa; skeletal characters | 12 | 4 | Marx & Fordyce (2015) |
| Mammalia: Eutheria: Primates | Lemuridae | 5767 bp from 120 taxa; 2 mt genes (*CYTB*, *ND4*) and 4 nuc genes (*ADORA*, *CNR*, *RAG-1*, *RAG-2*) | 421 chars from 85 taxa; cranial and long bone, dental, and transformed quantitative characters | 52 | 10 | Herrera & Dávalos (2016) |
| Mammalia: Eutheria: Rodentia | Abrotrichini | 4441 bp from 26 taxa; 1 mt gene (*CYTB*) and 5 nuc genes (*DMP1*, *IRBP*, *Adh*, *βfbg*, *Ins*) | 99 chars from 26 taxa; integument, skull, dental, postcranial, male genitalia, digestive system | 24 | 6 | Cañon et al. (2014); Teta et al. (2016) |

[a]These data sets were acquired directly from Keating et al. (2023); however, the original publication references and data set descriptions are included in this table.

[b]Abbreviations: cp = chloroplast, mt = mitochondrial, nuc = nuclear.

[c]Number of discrete morphological characters (chars) in the data set before pruning taxa and removing invariant sites.

[d]Number of taxa in the combined molecular and morphological data set, after pruning non-intersecting taxa.

**Supplementary Table S2.** Coefficient of variation (CoV) of root-to-tip distances estimated for molecular and morphological data sets. CoV was estimated by dividing the standard deviation of root-to-tip distances by the mean root-to-tip distance. Smaller values indicate lower degrees of rate variation across lineages. Root-to-tip distances were estimated from phylograms inferred using maximum-likelihood analysis.

| **Clade** | **Taxon** | **Coefficient of variation of root-to-tip distances** | |
|---|---|---|---|
| | | **Molecules** | **Morphology** |
| Alveolata | Avian Haemosporida | 0.4595 | 0.3228 |
| Plantae: Marchantiophyta | Marchantiidae | 0.3787 | 0.3127 |
| Plantae: Monilophyta | Marattiales | 0.3345 | 0.2811 |
| Plantae: Spermatophyta | Acrogymnospermae | 0.4343 | 0.4698 |
| Plantae: Monocotyledoneae | Arecaceae | 0.2878 | 0.4075 |
| Plantae: Monocotyledoneae | *Herminium* | 0.3012 | 0.3174 |
| Plantae: Eudicotyledoneae | Ericales | 0.2883 | 0.3044 |
| Plantae: Eudicotyledoneae | *Leptocereus* | 0.2201 | 0.3261 |
| Porifera | Hexactinellida | 0.1098 | 0.3218 |
| Platyhelminthes: Monogenea | Dactylogyridae | 0.2133 | 0.8095 |
| Platyhelminthes: Tricladida | Geoplaninae | 0.2155 | 0.4444 |
| Brachiopoda | Brachiopoda | 0.2588 | 0.4801 |
| Annelida: Polychaeta | Terebelliformia | 0.3816 | 0.6401 |
| Annelida: Polychaeta | Fabriciidaea | 0.269 | 0.3902 |
| Annelida: Polychaeta | Serpulidae | 0.2574 | 0.3688 |
| Arthropoda | Arthropoda | 0.3396 | 0.4899 |
| Arthropoda: Arachnida | Opiliones | 0.2304 | 0.2362 |
| Arthropoda: Arachnida: Araneae | Dionycha | 0.1872 | 0.3991 |
| Arthropoda: Arachnida: Araneae | Filistatidae | 0.1914 | 0.1156 |
| Arthropoda: Arachnida: Araneae | Nephilidae | 0.297 | 0.4916 |
| Arthropoda: Crustacea | Eumalacostraca | 0.8057 | 0.3308 |
| Arthropoda: Crustacea | Stomatopoda | 0.2622 | 0.4528 |
| Arthropoda: Insecta: Hemiptera | Heteroptera | 0.2473 | 0.2835 |
| Arthropoda: Insecta: Hemiptera | Reduviidae | 0.2129 | 0.3813 |
| Arthropoda: Insecta: Hemiptera | Coccoidea | 0.2957 | 0.4154 |
| Arthropoda: Insecta: Hemiptera | Coelidiinae | 0.0922 | 0.1874 |

| | | | |
|---|---|---|---|
| Arthropoda: Insecta: Hymenoptera | Hymenoptera | 0.266 | 0.4465 |
| Arthropoda: Insecta: Hymenoptera | Pimpliformes | 0.2856 | 0.3602 |
| Arthropoda: Insecta: Hymenoptera | *Polistes* | 0.4358 | 0.2344 |
| Arthropoda: Insecta: Coleoptera | Aleocharinae | 0.1744 | 0.1609 |
| Arthropoda: Insecta: Coleoptera | Entiminae | 0.4647 | 0.4054 |
| Arthropoda: Insecta: Coleoptera | Saprininae | 0.185 | 0.3654 |
| Arthropoda: Insecta: Neuroptera | Chrysopidae | 0.1818 | 0.1702 |
| Arthropoda: Insecta: Neuroptera | Hemerobiidae | 0.2914 | 0.3273 |
| Fishes: Chondrichthyes | Squaliformes | 0.2296 | 0.2992 |
| Fishes: Chondrichthyes | Scyliorhinidae | 0.29 | 0.3991 |
| Fishes: Osteichthyes | Tetradontiformes | 0.6079 | 0.4992 |
| Fishes: Osteichthyes | Catostomidae | 0.1704 | 0.2561 |
| Fishes: Osteichthyes | Anostomidae | 0.1939 | 0.5424 |
| Fishes: Osteichthyes | Gymnotiformes | 0.1916 | 0.3156 |
| Fishes: Osteichthyes | Loricariinae | 0.1652 | 0.842 |
| Reptilia: Squamata | Squamata | 0.319 | 0.8479 |
| Reptilia: Squamata | Iguania | 0.2462 | 0.7049 |
| Reptilia: Squamata | Serpentes | 0.29 | 0.3991 |
| Reptilia: Squamata | Crotalinae | 0.1759 | 0.2619 |
| Reptilia: Archosauria: Crocodylia | Crocodylia | 0.1266 | 0.5897 |
| Reptilia: Archosauria: Aves | Aves | 0.2697 | 0.2219 |
| Reptilia: Archosauria: Aves | Neoaves | 0.3884 | 0.3719 |
| Reptilia: Archosauria: Aves | Charadriiformes | 0.1109 | 0.3316 |
| Reptilia: Archosauria: Aves | Spheniscidae | 0.1005 | 0.2079 |
| Mammalia: Metatheria | Dasyuromorphia | 0.0728 | 0.2462 |
| Mammalia: Metatheria | Macropodidae | 0.095 | 0.4367 |
| Mammalia: Eutheria | Placentalia | 0.2741 | 0.4309 |
| Mammalia: Eutheria: Laurasiatheria | Phyllostomidae | 0.1417 | 0.688 |
| Mammalia: Eutheria: Laurasiatheria | Pinnipedia | 0.15 | 0.3036 |
| Mammalia: Eutheria: Laurasiatheria | Bovidae | 0.1157 | 0.4632 |
| Mammalia: Eutheria: Laurasiatheria | Neoceti | 0.2146 | 0.2759 |
| Mammalia: Eutheria: Laurasiatheria | Mysticetia | 0.1793 | 0.3455 |
| Mammalia: Eutheria: Primates | Lemuridae | 0.1374 | 0.2952 |
| Mammalia: Eutheria: Rodentia | Abrotrichini | 0.1037 | 0.6503 |

**Supplementary Table S3.** Details of assumption tests and number of contrasts used for independent sister-pair contrast analyses. Kendall's tau ($\tau$) and *p*-value are provided for each assumption test. Any trend ($p < 0.05$, bold font) indicates that assumptions are violated. Table does not show data sets that severely violated assumptions and so underwent a two-sample sign test.

| Group | Data type | Number of contrasts[a] | Assumption test 1 | | Assumption test 2 | |
|---|---|---|---|---|---|---|
| | | | Kendall's $\tau$ | *p*-value | Kendall's $\tau$ | *p*-value |
| Acrogymnospermae | Molecules | 5 | 0.2 | 0.82 | 0.2 | 0.82 |
| | Morphology | | –0.2 | 0.82 | 0 | 1.0 |
| Aleocharinae | Molecules | 13 | 0.077 | 0.77 | 0.077 | 0.77 |
| | Morphology | | –0.31 | 0.16 | –0.54 | **0.01** |
| Arecaceae | Molecules | 14 | 0.033 | 0.91 | 0.12 | 0.59 |
| | Morphology | | –0.011 | 1.0 | –0.033 | 0.91 |
| Arthropoda | Molecules | 8 | 0.5 | 0.11 | 0.5 | 0.11 |
| | Morphology | | 0.14 | 0.72 | –0.071 | 0.90 |
| Aves | Molecules | 24 | –0.25 | 0.097 | 0.051 | 0.75 |
| | Morphology | | –0.022 | 0.90 | 0.16 | 0.29 |
| Avian Haemosporida | Molecules | 15 | –0.067 | 0.77 | –0.12 | 0.56 |
| | Morphology | | –0.30 | 0.14 | –0.56 | **0.0029** |
| Bovidae | Molecules | 12 | –0.39 | 0.086 | –0.36 | 0.12 |
| | Morphology | | –0.30 | 0.20 | –0.03 | 0.95 |
| Catostomidae | Molecules | 13 | –0.33 | 0.13 | –0.33 | 0.13 |
| | Morphology | | –0.18 | 0.44 | 0.077 | 0.77 |
| Chrysopidae | Molecules | 13 | –0.15 | 0.51 | –0.15 | 0.51 |
| | Morphology | | –0.21 | 0.37 | –0.41 | 0.057 |
| Crotalinae | Molecules | 14 | –0.011 | 1.0 | 0.25 | 0.23 |
| | Morphology | | 0.12 | 0.59 | –0.23 | 0.28 |
| Dionycha | Molecules | 30 | –0.20 | 0.12 | –0.15 | 0.26 |
| | Morphology | | –0.12 | 0.38 | –0.044 | 0.75 |
| Eumalacostraca | Molecules | 4 | –0.67 | 0.33 | 0.33 | 0.75 |
| | Morphology | | 0 | 1.0 | 0 | 1.0 |
| Filistatidae | Molecules | 7 | –0.24 | 0.56 | 0.24 | 0.56 |
| | Morphology | | 0.047 | 1.0 | 0.14 | 0.77 |
| Hexactinellida | Molecules | 8 | –0.14 | 0.72 | 0 | 1.0 |
| | Morphology | | –0.071 | 0.90 | –0.43 | 0.18 |
| Hymenoptera | Molecules | 10 | –0.24 | 0.38 | –0.11 | 0.73 |

| | | | | | | |
|---|---|---|---|---|---|---|
| | Morphology | | –0.42 | 0.11 | –0.33 | 0.22 |
| Iguania | Molecules | 7 | –0.42 | 0.24 | –0.14 | 0.77 |
| | Morphology | | –0.52 | 0.14 | –0.24 | 0.56 |
| Lemuridae | Molecules | 6 | –0.6 | 0.14 | –0.33 | 0.47 |
| | Morphology | | –0.2 | 0.72 | –0.33 | 0.47 |
| Loricariinae | Molecules | 7 | –0.24 | 0.56 | –0.24 | 0.56 |
| | Morphology | | –0.33 | 0.38 | 0.14 | 0.77 |
| Macropodidae | Molecules | 5 | –0.4 | 0.48 | 0.33 | 0.47 |
| | Morphology | | –0.47 | 0.27 | 0.067 | 1.0 |
| Marchantiidae | Molecules | 7 | 0.14 | 0.77 | –0.24 | 0.56 |
| | Morphology | | 0.048 | 1.0 | 0.14 | 0.77 |
| Mysticeti | Molecules | 4 | 0 | 1.0 | 0.33 | 0.75 |
| | Morphology | | –0.33 | 0.75 | 0.33 | 0.75 |
| Neoaves | Molecules | 8 | –0.29 | 0.40 | –0.21 | 0.55 |
| | Morphology | | 0 | 1.0 | 0.21 | 0.55 |
| Neoceti | Molecules | 7 | –0.048 | 1.0 | 0.048 | 1.0 |
| | Morphology | | –0.33 | 0.38 | –0.14 | 0.77 |
| Nephilidae | Molecules | 11 | 0.018 | 1.0 | 0.055 | 0.88 |
| | Morphology | | –0.31 | 0.22 | –0.38 | 0.12 |
| Phyllostomidae | Molecules | 10 | –0.16 | 0.60 | 0.022 | 1.0 |
| | Morphology | | –0.38 | 0.16 | –0.24 | 0.38 |
| Pimpliformes | Molecules | 23 | –0.18 | 0.25 | –0.051 | 0.75 |
| | Morphology | | 0.28 | 0.88 | –0.41 | **0.0061** |
| Placentalia | Molecules | 13 | 0.21 | 0.37 | 0.41 | 0.057 |
| | Morphology | | –0.28 | 0.20 | –0.10 | 0.68 |
| *Polistes* | Molecules | 11 | –0.091 | 0.76 | 0.16 | 0.54 |
| | Morphology | | 0.055 | 0.88 | –0.091 | 0.76 |
| Reduviidae | Molecules | 7 | –0.33 | 0.38 | –0.33 | 0.38 |
| | Morphology | | 0.33 | 0.38 | 0.52 | 0.14 |
| Saprininae | Molecules | 16 | –0.5 | 0.82 | 0.017 | 0.96 |
| | Morphology | | –0.017 | 0.96 | –0.13 | 0.51 |
| Serpentes | Molecules | 13 | –0.28 | 0.20 | 0.051 | 0.86 |
| | Morphology | | 0.077 | 0.77 | 0.21 | 0.37 |
| Squamata | Molecules | 49 | –0.16 | 0.11 | –0.049 | 0.63 |
| | Morphology | | –0.10 | 0.30 | 0 | 1.0 |
| Terebelliformia | Molecules | 20 | 0.11 | 0.54 | 0.22 | 0.19 |
| | Morphology | | 0.14 | 0.42 | –0.2 | 0.23 |

[a]Number of contrasts after removing any unsuitable contrasts. This is the number of contrasts used for all assumption tests and for the final comparison of molecular and morphological contrasts.

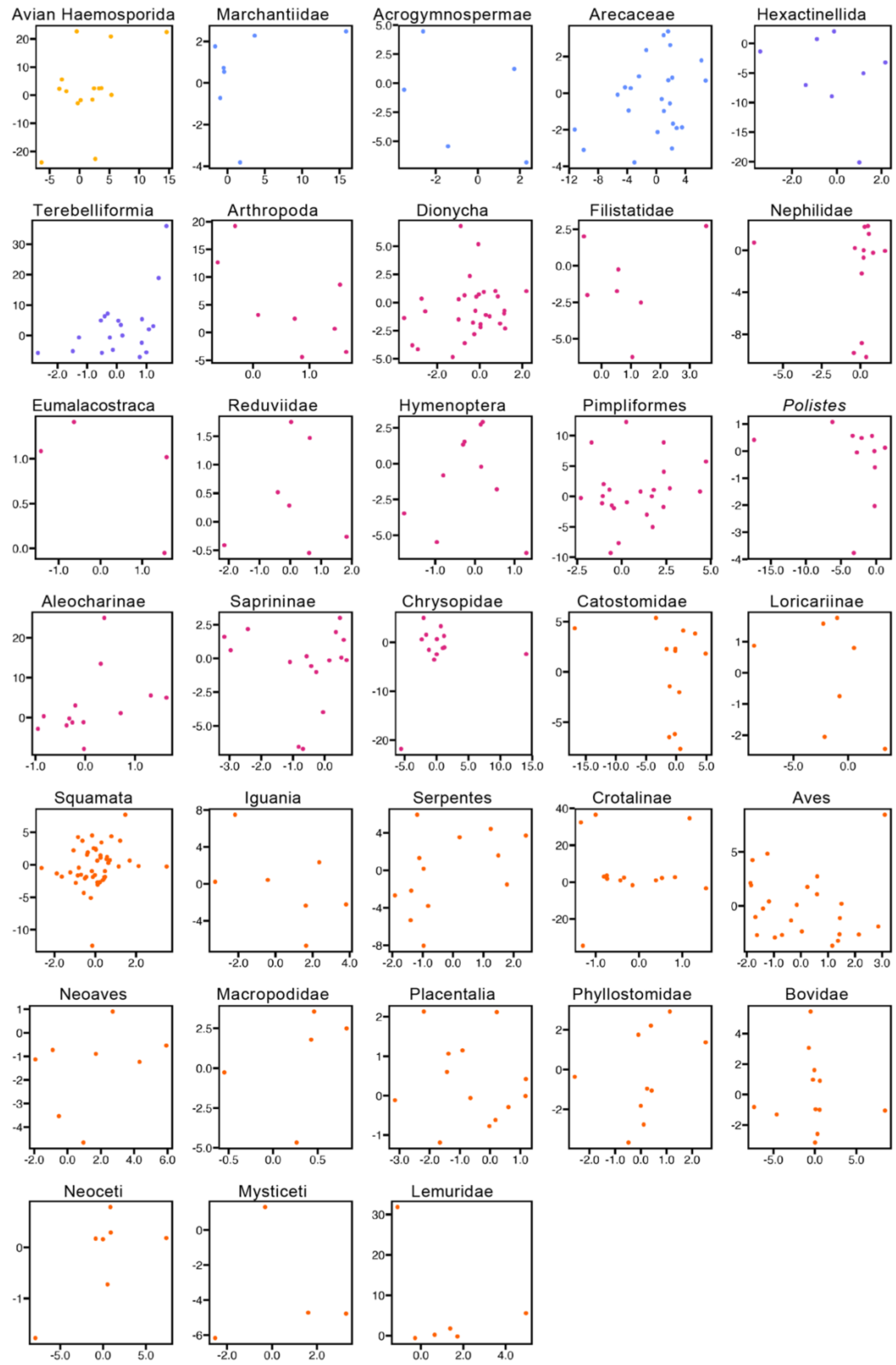


**Supplementary Figure S1.** Plots of standardized, log-transformed sister-pair contrasts of molecular branch lengths and morphological branch lengths. Plots are shown for the 33 data sets that passed the assumption tests. Correlation coefficients (*rho*) and *p*-values are given in figure 1.
